# Bacteriophage adherence to mucus mediates preventive protection against pathogenic bacteria

**DOI:** 10.1101/592097

**Authors:** Gabriel MF Almeida, Elina Laanto, Roghaieh Ashrafi, Lotta-Riina Sundberg

## Abstract

Metazoan mucosal surfaces are major interfaces between the organism and environment. These surfaces have been proposed to host bacteriophages in a symbiotic relationship with metazoans. Considering the so far poorly understood phage–mucus interaction and its role in ecological interactions and for mucosal bacterial infections, empirical evidence and model systems need to be established. Here, using the fish pathogen *Flavobacterium columnare* and rainbow trout (*Oncorhynchus mykiss*), we show that phages infecting the pathogen are capable of binding to primary mucus layers and protecting fish from infections. Furthermore, exposure to mucus changes the bacterial phenotype by increasing bacterial virulence and susceptibility to phage infections. Tests using other phage–bacterium pairs suggest that the relevance of mucus for bacteria and phages may be widespread in the biosphere. Therefore, interactions of bacteria and phages inside the mucus environment may be important for disease and evolution, and this phenomenon has significant potential to be exploited for preventive phage therapy approaches.

## Main text

Mucus is an essential component of the innate immune system, serving as a selective barrier between metazoans and their environments^**1**^. It is a complex, viscoelastic secretion that, in addition to protecting the host, provides a habitat for countless microbes from all three domains of life, as well as viruses. Mucins, the main components of the mucosal matrix, form a polymer-based hydrogel with lubricant and protective properties. The mucin concentration of the mucus affects the mesh size of the matrix, which controls diffusion through it and provides a first line of defence against pathogens^**2**^. Most infections start from the mucosal surfaces, and with over 200 known microorganisms capable of invading the mucus layers, mucosal infections result in mortality and morbidity, exhibiting clinical and economical importance worldwide^**3,4**^. Considering the heterogeneous composition of mucosal surfaces, there is a significant gap in knowledge on how its components interact during mucosal infections. Previous research has mainly focussed on the pathogen, on the host immune response *in vitro* or the effect of the commensal microbiome for mucosal homeostasis^**5-7**^. Moreover, in 2013, another layer of complexity was added to the already convoluted system when the bacteriophage adherence to mucus (BAM) model was proposed^**8**^. This model, based on indirect evidence and *in vitro* testing, proposes an important and so far overlooked symbiosis between metazoans and bacteriophages (phages): Phages would concentrate on mucosal surfaces by weak interactions with mucins, creating a ubiquitous non-host-derived immunity against bacterial invaders during the mucus colonisation process. Phage structural proteins possessing immunoglobulin (Ig)-like folds were proposed to be mediators for phage interaction with mucins. A bioinformatic analysis of 246 double stranded DNA tailed-phage genomes revealed that roughly 25% have proteins with Ig-like folds, all related to the viral structure^**9**^. Indeed, the virus abundance in the surface mucus layer of eels has been shown to be higher compared with surrounding water; the sub-diffusive motion of phages on mucus has been better understood; and the spatial structuring of mucosal surfaces has been speculated to have a role in phage replication strategies^**10-12**^. However, the relevance of phage interaction with metazoan mucus for bacterial infections has not been shown in a natural infection model.

Fish are naturally covered by mucus layers, and thus, logical model organisms for studying phage–mucus interactions and their consequent influences on bacterial infections, especially those that affect mucosal surfaces. *Flavobacterium columnare* (phylum Bacteroidetes, family Flavobacteriaceae), the causative agent of columnaris disease, is a major epidermal pathogen in freshwater aquaculture worldwide and responsible for annual losses in the magnitude of millions of dollars^**13,14**^. Previous studies have indicated that mucus has a positive effect on *F. columnare* growth, chemotaxis, motility gene upregulation and protease production, demonstrating that interaction with the mucosal surfaces is important for the bacterial pathogenesis^**15-19**^. Treatment of columnaris disease is based on antibiotics, which lead to antibiotic residues in fish and antibiotic leakage to the environment. Antibiotic use may also select for antibiotic-resistant bacteria^**20**^. Phages capable of infecting *F. columnare* have already been described and shown to be efficient as phage therapy agents in laboratory conditions^**21**^. However, so far, no study has focussed on the interaction between *F. columnare* and phages on the mucosal environment.

Considering that *F. columnare* infections represent a relevant model for studying bacterial mucosal diseases, and in the context of columnaris disease, the fish skin mucosa could benefit from phage protection as predicted by the BAM model, we decided to study how phage–host interactions occur in this system. First, we verified that a phage specific to *F. columnare* remains associated with the fish skin mucosa for as long as 7 days after a single exposure. Next, we showed that phage pre-treatment protects fish from a subsequent *F. columnare* infection, in accordance with the BAM model. Then, to better understand the interactions of phage and bacteria in mucus, we devised a simulated mucus environment to grow the bacterium and discovered that it influences bacterial virulence and phage growth. Finally, we tested other phage–bacteria pairs, discovering that bacterial physiology modification by mucin exposure and exploitation of the mucin-generated state by phages is not limited to *F. columnare* and may be widespread in the biosphere. Our data represent the first evidence of phage adherence to mucus relevance in a natural mucosal infection system. Phage–mucus interactions open promising possibilities for the development of preventive phage therapy approaches for mucosal diseases caused by bacteria in farmed animals and humans.

## Results

### Ig-like domains in phage structural proteins support the suggestion of interaction mechanisms with mucus

Considering the importance of Ig-like domains for the interaction of phages and mucus, we tested whether distinct phages from our collection would contain these domains in their structural proteins. *In silico* re-annotation of the FCL-2^**21**^ genome using HHPred revealed one open reading frame (orf) (orf17, length 114 amino acids, accession YP_009140518) homologous to an N-terminal domain of orf48 in *Lactococcus* phage TP901-1 (probability 98.94, E-value 4.5 E-10). The same result was achieved using Phyre2 for homology detection (confidence 86.5%, coverage 53%). The orf48 of TP901-1 has been experimentally shown to encode a baseplate protein with an Ig fold^**22**^. In the FCL-2 genome, orf17 is located between a putative peptidase and two hypothetical proteins that are then followed by a putative portal protein and capsid proteins, suggesting that there could be structural function for its protein. The genome sequencing of *Aeromonas* sp. podovirus V46 (unpublished) revealed a capsid protein that was highly similar to phage P22, including the telokin-like Ig domain. In addition, the Phyre2 homology recognition resulted in 100% confidence on the probability that the sequences are homologous. The two *F. psychrophilum* phages and *Flavobacterium* sp phage FL-1 used in the study did not show apparent homology to Ig-like folds in their predicted orfs and neither did the structural proteins of the two tailless phages used, FLiP and PRD1

### The skin mucosa of rainbow trout retains phages for one week

To verify phage FCL-2 retention by primary mucus *in vivo*, we exposed rainbow trout individuals to FCL-2 and kept the fish in flow-through aquaria with a water flow of 860 ml/min. The daily phage titres, in plaque forming unities (pfu) ml^−1^, were obtained from the water and fish skin mucus. Phage titres in water decreased rapidly, and at some timepoints, no phages were detected. However, phages were found on the mucus consistently for up to 7 days after phage exposure, despite the water flow and mucus shedding (**Fig. 1**). No phages were detected on water or mucus 15 days after exposure (data not shown).

**Figure 1:**
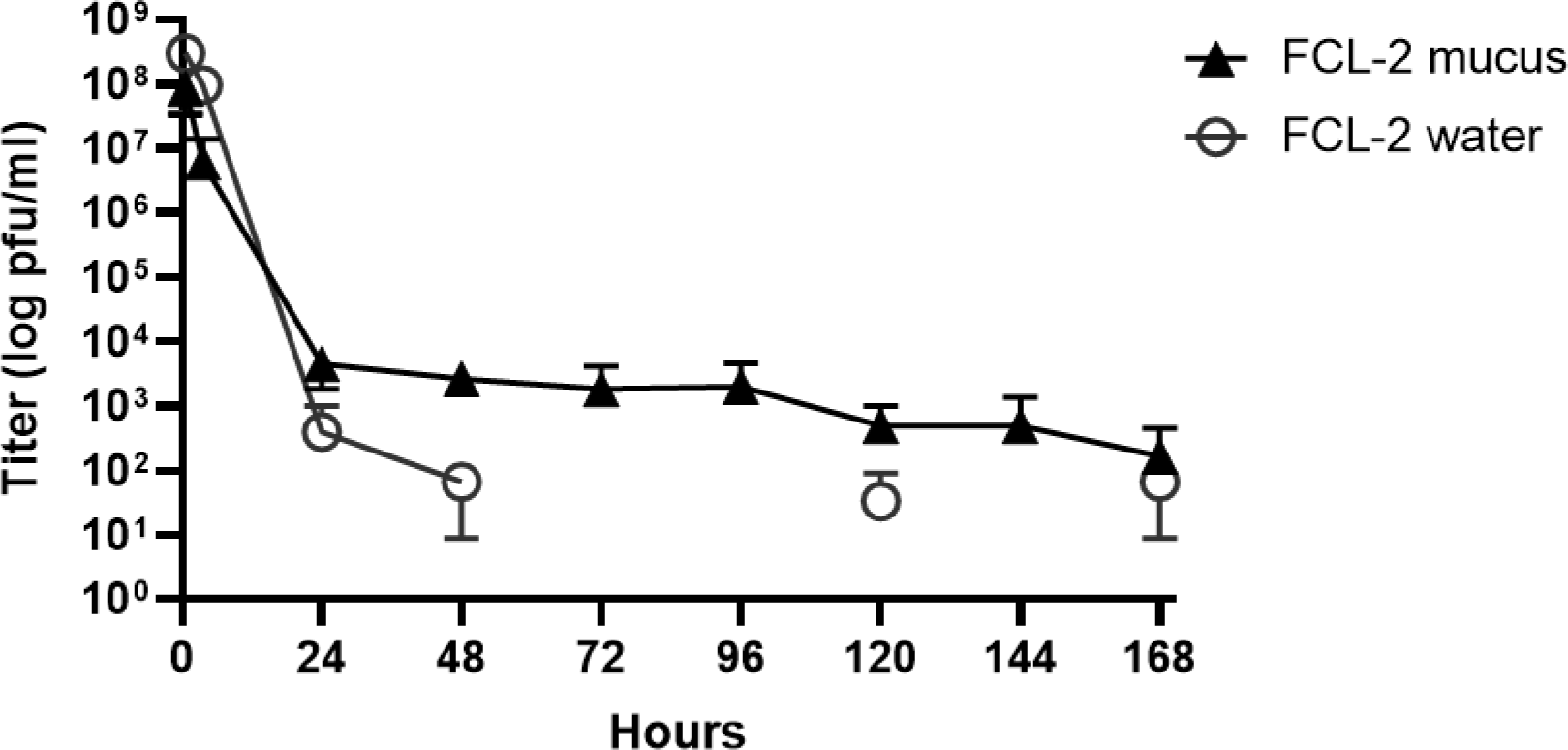
Phages persist on the rainbow trout skin mucosa for up to 1 week after a single exposure. Phage FCL-2 titres in water (grey open circles) and fish mucus (black triangles) over time in a flow-through aquarium. Fish were exposed to the phage only once, at time 0, and water flow was opened after 3.5 h. Each data point represents the average of three individual fish or three independent water samples and their standard deviation.

### Pre-treatment with FCL-2 phage protects fish from columnaris disease

Phage retention by fish skin mucus demonstrates that the dynamics predicted by the BAM model can be important for the outcome of columnaris disease. To test this possibility, we exposed rainbow trout to FCL-2 phage for 1, 3 or 7 days and then infected the fish with *F. columnare*. For exploring the influence of a co-occurring phage on FCL-2 mucus adhesion and fish protection, we also exposed the fish to a mixture of FCL-2 and T4 for 1 or 7 days before infection. Right before infection, the phage particles were quantified by titration from fish mucus and the water in which the fish were kept to verify the phage retention in the mucus. FCL-2 was detected in the mucus of fishes from all the pre-exposed groups, but it was only found in the water collected from the aquaria in which exposure occurred 1 or 3 days before infection (**Fig. 2a** and **2b**). T4 was detected on mucus from fish exposed for 1 and 7 days, but it was only recovered from the water collected from the aquaria in which the exposure occurred 1 day before (**Fig. 2b**). The T4 presence did not affect FCL-2 adhesion to mucus, and both phages were recovered at similar titres from the fish. Exposing the fish to FCL-2 alone or FCL-2 mixed with T4 before the *F. columnare* challenge resulted in a delay of columnaris disease and increased fish survival. Fish pre-treated with FCL-2 alone for 1 and 3 days survived longer than controls (*p*-values of 0.0398 and 0.0013, respectively; **Fig. 2c**). When combined with T4, FCL-2 could protect fish in the 1- and 7-day pre-treatment groups (*p* < 0.0001 and 0.0003, respectively; **Fig. 2d**). Interestingly, the presence of T4 enhanced the protective effect of FCL-2, as fishes exposed 7 days before infection to T4 and FCL-2 had higher survival rates than fishes exposed only to FCL-2 in the same timeframe (*p* = 0.0008).

**Figure 2:**
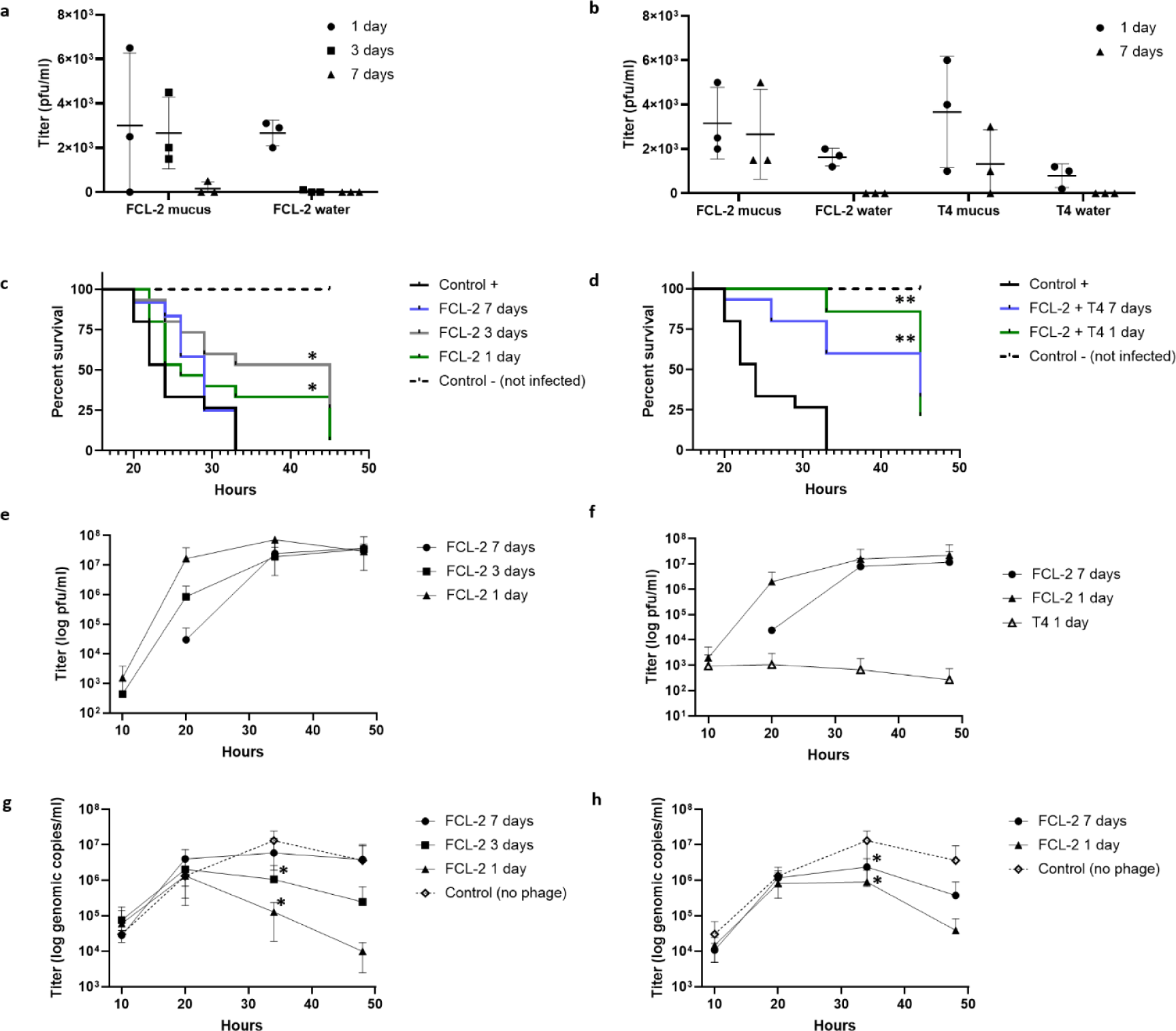
Phage retention on mucus allows for the prophylactic use of phages against bacterial infections. Fish were pre-treated with FCL-2 alone (1, 3 or 7 days) or with FCL-2 and T4 (1 or 7 days) before infection with *Flavobacterium columnare*. (**a**) FCL-2 phage titres in fish mucus and aquarium water (before infection) from FCL-2-only groups. (**b**) FCL-2 and T4 titres from fish mucus and aquarium water (before infection) in the FCL-2 combined with T4 groups. (**c**) Fish survival in the FCL-2 only pre-treated groups. (**d**) Fish survival in the FCL-2 combined with T4 pre-treated groups. (**e**) FCL-2 titres in water after infection in the FCL-2-only groups. (**f**) FCL-2 and T4 titres in water after infection on the FCL-2 combined with T4 groups. (**g**) *F. columnare* quantity (genomic copies) in water after infection in the FCL-2 only pre-treated groups. (**h**) *F. columnare* quantity (genomic copies) in water after infection in the FCL-2 combined with T4 pre-treated groups. Each data point represents one individual fish or water sample and the standard deviations in **a** and **b**. In **c** and **d**, the number of fishes used was 15 per group except in the negative control (*n* = 16) and the group exposed to FCL-2 alone for 7 days (*n* = 12). In **e**, **f**, **g** and **h**, the data points represent the average of triplicates and their standard deviation. The log-rank (Mantel–Cox) test was used for evaluating the survival curves in **c** and **d**. Unpaired *t*-tests were used for comparing the controls and tested conditions in **g** and **h** (\**p* < 0.05, \*\**p* < 0.001).

The FCL-2 titres in the water increased over time, peaking around 35 h after exposure to *F. columnare*. The phage numbers were highest in the water of fish pre-treated for 1 day prior to infection, followed by fish pre-treated for 3 days and then 7 days before infection (**Fig. 2e** and **2f**). T4 was detected at low levels in only one aquarium from the 1-day pre-treatment group at all timepoints (**Fig. 2f**). *F. columnare* genome copies from the water were measured by quantitative PCR, and they were found to be inversely proportional to the FCL-2 titres (**Fig. 2g** and **2h**). When compared with the controls, the reduction of bacterial genome copies was significant at 35 h after infection in the groups pre-treated with FCL-2 alone at 1 and 3 days before infection (*p* = 0.005 and 0.002, respectively) and groups pre-treated with FCL-2 and T4 at 1 and 7 days before infection (*p* = 0.004 and 0.009, respectively).

### Exposure to a simulated mucosal environment influences *F. columnare* growth characteristics and virulence factors

The data above revealed that the mucus environment affected both *F. columnare* and phages, suggesting that the mucus surface is a relevant site for interactions. For better understanding phage–host interactions in the presence of mucus, *F. columnare* strain B185 was grown in control conditions (0.5× Shieh) or 0.5× Shieh supplemented with purified porcine mucin, which was used for reproducibility purposes. One peculiarity of *F. columnare* cultures containing mucin is the formation of a thick biofilm ring on the liquid– air interface and smaller biofilm strings spread over the tube walls (**Fig. 3a**). Helium ion microscopy imaging of the mucin-generated biofilm revealed that it was composed of large masses of organised cells, growing as independent structures on the colonised surfaces (**Fig. 3b–3d**). No cells were visualised in agar blocks added to control cultures (data not shown), meaning that there was no adhesion or the cells detached during sample preparation. In addition biofilm formation, mucin cultures also induced positive chemotaxis, with an average relative chemotaxis response (RCR) of 12 to 0.1% mucin and 73 to 1% mucin containing Shieh medium (**Fig. 3e**). This can be taken as a strong chemotactic response, since some authors consider an RCR over 2 to be significant^**23**^. *F. columnare* spread on agar was also improved by mucin presence (**Fig. 3f**; *p* < 0.0001 comparing controls with agar supplemented with 0.2% or 1% mucin). Protease secretion was more prominent on agar plates containing mucin and skimmed milk (**Fig. 3g**). A fish infection experiment revealed that a lower dose of *F. columnare* grown in mucin (11.8 times fewer cells) had the same virulence as a higher dose of cells from control cultures (*p* = 0.3137, **Fig. 3h**, black and grey lines). In addition, the cells from mucin cultures killed fish more efficiently than a dose 2.3 times higher from control cultures (*p* = 0.0314, **Fig. 3h**, black and blue lines). The survival of fish exposed to control cells was dose dependent (*p* = 0.0285, **Fig. 3h**, grey and blue lines).

**Figure 3:**
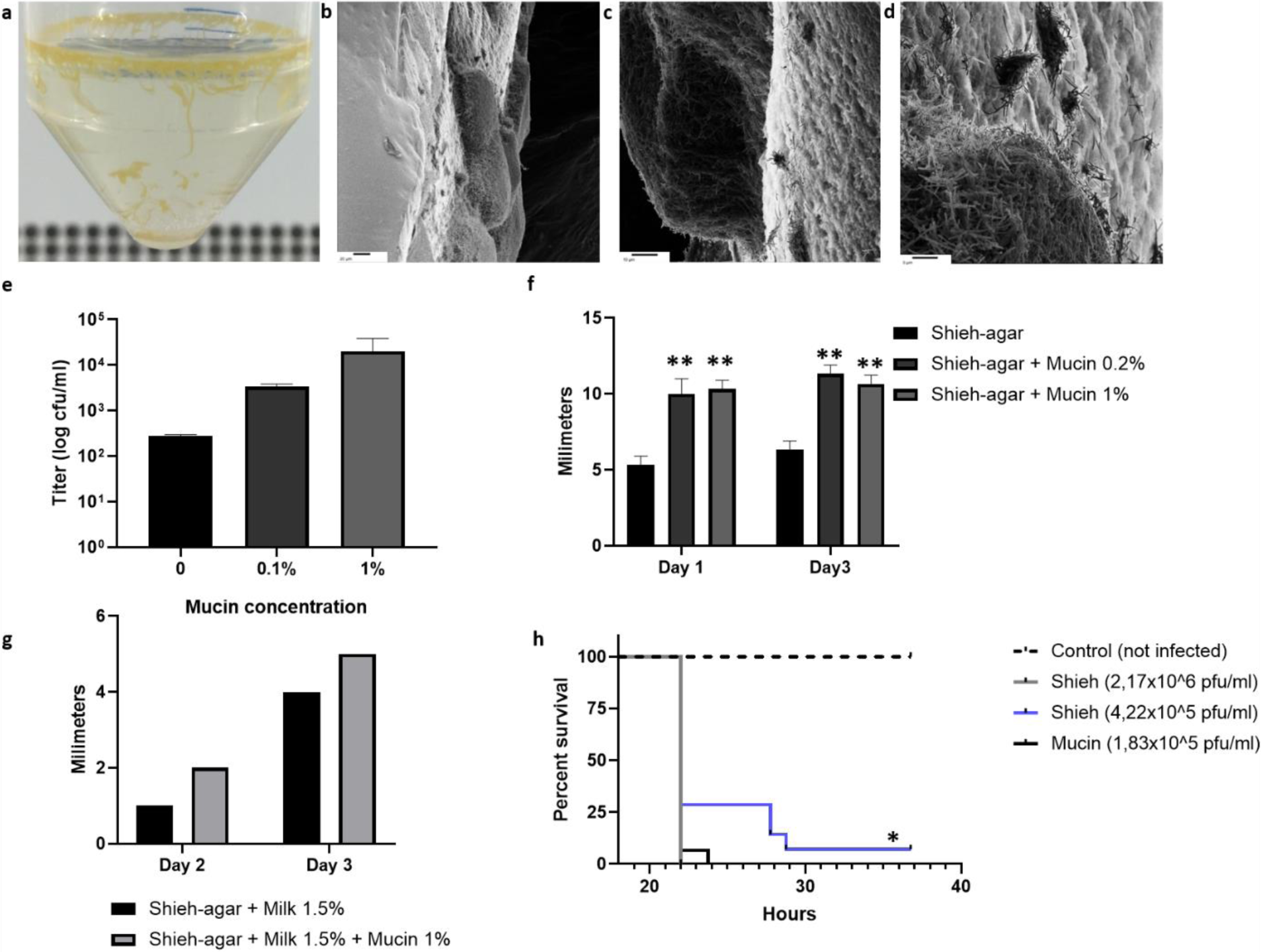
Exposure to a simulated mucosal environment increases *Flavobacterium columnare* virulence. (**a**) Macroscopic view of a 22-h-old culture of *F. columnare* in mucin containing media. Note the thick biofilm ring on the liquid–air interface, biofilm strings and clearer supernatant. (**b**–**d**) Helium ion microscopy images of the *F. columnare* mucin-induced biofilm colonising a block of agar. (**e**) *F. columnare* chemotaxis to mucin containing Shieh media. (**f**) *F. columnare* spread (growth size) on solid Shieh media containing mucin. (**g**) *F. columnare* protease halo formed on skimmed milk containing plates, in the presence or absence of mucin. (**h**) Fish survival from a virulence test *in vivo* comparing *F. columnare* grown in 0.5× Shieh media to *F. columnare* grown in 0.5× Shieh supplemented with 0.1% mucin. Each data point represents the average of duplicates their standard deviation in **e**, which is representative of two independent experiments. In **f** and **g**, the data points represents the average of triplicates and their standard deviation. In **h**, the total number of fishes used was 15 per group except mucin 1.83 × 10^5^ (*n* = 14). Unpaired *t*-tests were used for comparing controls and tested conditions on **f**. The log-rank (Mantel–Cox) test was used to evaluate survival curves in **h** (\**p* < 0.05, \*\**p* < 0.001).

The influence of mucin on bacterial growth characteristics was seen for all *F. columnare* strains tested (B067, B185, B245, B350, B407, B420, B480, B537, G1, H2, JIP39/87 and JIP44/87), covering genotypes A/C/E/G/H and two reference strains. It was assessed by the appearance of a biofilm ring on the mucin-containing cultures (**Table 1**).

**Table 1:** List of bacterial strains and phages used in this study, and their behavior concerning biofilm formation and phage growth on culture media containing purified mucin.

| Bacterial sample | Culture aspect changes on mucin cultures | Tested phages | Increased phage yield on mucin cultures |
| --- | --- | --- | --- |
| *Aeromonas salmonicida*<sup>1</sup> | No | None | - |
| *Aeromonas sp B135*<sup>2</sup> | Yes (biofilm) | V46<sup>2</sup> | Yes |
| *Aeromonas sp B158*<sup>2</sup> | Yes (biofilm) | V61<sup>2</sup> | Yes<sup>o</sup> |
| *Chryseobacterium indologenes*<sup>2</sup> | No | FCL-2<sup>2</sup> * | No |
| *Escherichia coli* DSM613<sup>3</sup> | No | T4 DSM4505<sup>3</sup> | No |
| *F. johnsoniae* UW101<sup>4</sup> | No | None | - |
| *Flavobacterium columnare* B067<sup>2</sup> | Yes (biofilm) | FCL-2<sup>2</sup> * | No |
| *Flavobacterium columnare* B185<sup>2</sup> | Yes (biofilm) | FCL-2<sup>2</sup> , FCOV-F46<sup>2</sup> , FCOV-F54<sup>2</sup> , FCOV-F55<sup>2</sup> | Yes |
| *Flavobacterium columnare* B245<sup>2</sup> | Yes (biofilm) | FCL-2<sup>2</sup> *, V156<sup>2</sup> | Yes for V156 |
| *Flavobacterium columnare* B350<sup>2</sup> | Yes (biofilm) | FCL-2<sup>2</sup> * | No |
| *Flavobacterium columnare* B407<sup>2</sup> | Yes (biofilm) | FCL-2<sup>2</sup> | Yes |
| *Flavobacterium columnare* B420<sup>2</sup> | Yes (biofilm) | FCL-2<sup>2</sup> | Yes |
| *Flavobacterium columnare* B480<sup>2</sup> | Yes (biofilm) | FCL-2<sup>2</sup> * | No |
| *Flavobacterium columnare* B537<sup>2</sup> | Yes (biofilm) | FCOV-F47<sup>2</sup> , FCOV-F49<sup>2</sup> , FCOV-F50<sup>2</sup> , FCOV-F51<sup>2</sup> , FCOV-F52<sup>2</sup> , FCOV-F59<sup>2</sup> , FCOV-F60<sup>2</sup> , FCOV-F61<sup>2</sup> , FCOV-F62<sup>2</sup> | Yes |
| *Flavobacterium columnare*<br>G1<sup>2</sup> | Yes (biofilm) | FCL-2<sup>2</sup> | Yes |
| *Flavobacterium columnare*<br>H2<sup>2</sup> | Yes (biofilm) | FCL-2<sup>2</sup> * | Yes |
| *Flavobacterium columnare*<br>JIP39/87<sup>5</sup> | Yes (biofilm) | FCL-2<sup>2</sup> *, V156<sup>2</sup> * | No |
| *Flavobacterium columnare*<br>JIP44/87<sup>5</sup> | Yes (biofilm) | FCL-2<sup>2</sup> *, V156<sup>2</sup> * | No |
| *Flavobacterium psychrophilum* 950106-1/1<sup>6</sup> | No | FPV4<sup>6</sup> , FPV9<sup>6</sup> | Negative effect<br>for both with<br>0.1% mucin |
| *Flavobacterium sp* B183<sup>2</sup> | No | FL-1<sup>2</sup> , FCL-2<sup>2</sup> * | No |
| *Flavobacterium sp* B330<sup>2</sup> | No | FLiP<sup>2</sup> | No |
| *Flavobacterium sp* strains<br>B28<sup>2</sup>, B167<sup>2</sup>, B222<sup>2</sup>, B225<sup>2</sup>,<br>B257<sup>2</sup> | No | None | - |
| *Pseudomonas fluorescens*<sup>1</sup> | No | None | - |
| *Salmonella enterica* Serovar<br>Typhimurium DS88<sup>2</sup> | No | PRD1<sup>2</sup> | No |
| *Yersinia ruckeri*<sup>1</sup> | No | None | - |

Origin of the samples: <sup>1</sup> Acquired from ATCC. <sup>2</sup> Our own collection (**Laanto et al 2017**<sup>47</sup> and unpublished data). <sup>3</sup> Acquired from DSMZ. <sup>4</sup> Kindly donated by Professor Mark McBride (University of Wisconsin-Milwaukee, USA). <sup>5</sup> Kindly donated by Professor Jean-Francois Bernardet (INRA, France). <sup>6</sup> Kindly donated by Professor Mathias Middelboe (University of Copenhagen, Denmark).
An \* after a phage name means that it was used to infect a non-susceptible host. ° Although a clear trend was verified, results could not be quantified due to small and turbid plaques.

### Exposure to a simulated mucosal environment makes *F. columnare* susceptible to phage infections

Since mucin exposure changes the *F. columnare* growth characteristics, we also considered whether the presence of mucin or mucus affects phage infections. To accomplish this, *F. columnare* strain B185 grown in control conditions (0.5× Shieh) or in 0.5× Shieh supplemented with primary fish mucus or purified porcine mucin was infected with phage FCL-2, and phage numbers were measured by titration at different time points. The culture aspect in primary mucus cultures was similar to that in mucin-containing cultures (biofilm formation). While the phage yield was negative in control cultures and cultures containing low mucus and mucin concentrations, it was hundreds- to thousands-fold higher than the initial inoculum at the higher doses (**Fig. 4a** and **4b**). When compared with controls, a significant increase in phage titres was detected 24 and 48 h after infection in media containing 13.25 mg ml^−1^ of mucus (*p* < 0.0001), 0.1% mucin (*p* = 0.008 and 0.00003, respectively) and 1% mucin (*p* = 0.007 and 0.00001, respectively).

**Figure 4:**
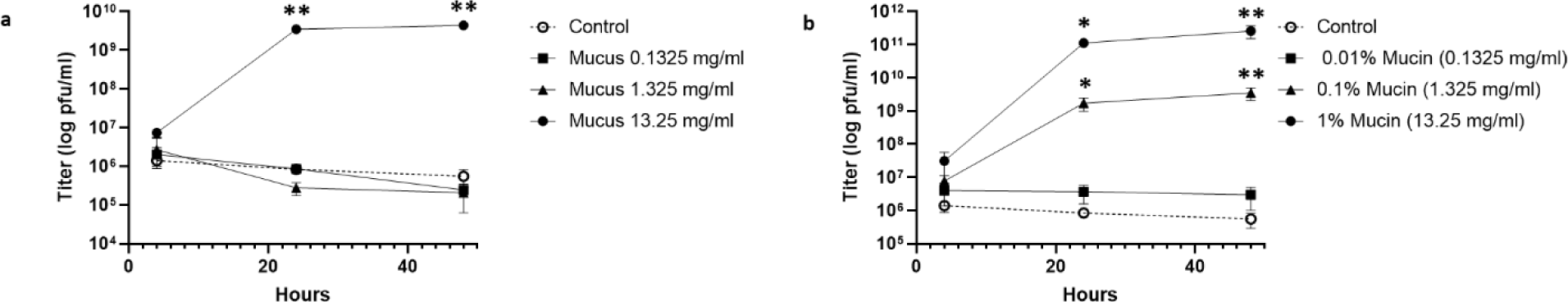
Exposure to simulated mucosal environments makes *Flavobacterium columnare* susceptible to phage infection. (**a**) FCL-2 titres over time on 0.5× Shieh cultures supplemented with primary rainbow trout mucus. (**b**) FCL-2 titres over time on 0.5× Shieh cultures supplemented with purified porcine mucin. Each data point represents the average of triplicates and their standard deviation. Unpaired *t*-tests were used for comparing the controls and tested conditions (\**p* < 0.05, \*\**p* < 0.001).

For elucidating the role of mucin-induced biofilms in phage infections, mucin cultures were prepared and had the biofilm and planktonic cells divided before infection with FCL-2. It was possible to see that phage replication was efficient in cultures containing the planktonic cells from mucin cultures and not in the controls or mucin-generated biofilm alone (**Supplementary Fig. 1a**). Phage production, measured by dividing the total yield by the inoculum, was negative in the controls and higher in cultures containing only planktonic cells than in cultures containing these cells and the biofilm together. For evaluating the phage effect on biofilm spread, biofilm baits colonised by *F. columnare* in the presence of mucin were transferred to fresh media containing mucin and clean baits. This qualitative analysis revealed that the phage presence impairs the ability of *F. columnare* grown in mucin to spread as biofilm to surfaces, but it does not kill biofilm formed before the addition of the phages (**Supplementary Fig. 1b**). The reduction of biofilm colonisation on the baits and formation of the typical biofilm ring on the mucin culture was phage dose dependent.

Due to the fast biofilm formation and strong biofilm structure, our attempts to titrate *F. columnare* cells grown in mucin returned variable results, since it is impossible to properly homogenise the cultures before making serial dilutions, optical density measurements or nucleic acid extractions. Diluting and plating only the plankton containing supernatant is possible, but this also results in variations in the final titres. However, from the plankton platings, it is clear that mucin exposure leads to the appearance of a novel colony morphotype. This is larger and does not attach as firmly to the agar as normal rhizoid colonies do (**Supplementary Fig. 2a**). To understand the duration of the mucin effect on cells, we followed the appearance of this novel colony morphotype and phage susceptibility on liquid over four passages of the original control and mucin cultures. The increase in phage titres was taken as a measure of phage susceptibility, while the new colony morphotype was used as a marker for the cell changes elicited by mucin exposure. After an initial exposure to mucin, the cultures were serially passaged daily for 3 days on media without mucin. The same was done for control cultures. Each passage was used for plating planktonic cells, and phage susceptibility was evaluated by infecting the cultures and subsequent quantification of phages by titration after 24 h. Increased phage susceptibility was only verified on the original cultures containing mucin (*p* = 0.0003) and disappeared in later passages (**Supplementary Fig. 2b**). However, the modified rhizoid colonies were evident in all passages of the mucin cultures, regardless of phage presence or absence. These colonies were not present in the control cultures and represented 0.07 ± 0.049 to 3.71 ± 1.31% of the total colonies in plankton from mucin cultures (**Supplementary Fig. 2c**). Biofilm formation ceased on the first passage, in which only clumps of cells were observed. From passage 2 onwards, all the cultures had the same turbid appearance.

Mucin exposure had a positive effect for the FCL-2 yield when hosts G1, B407, B420 (G genotypes) and H2 (H genotype) were used. Host H2 is resistant to FCL-2 infections in control cultures, and an effect for the phage host range was seen only in the presence of mucin. However, this was not a broad effect, since no productive infections were observed on hosts B067, B245, B350, JIP39/87 or JIP44/87 (*F. columnare* genotypes A, C and E and two reference strains; **Supplementary Fig. 3a**). Mucin cultures were also beneficial for other *F. columnare* phages, where 13 phages capable of infecting strains from genotypes C or G from our collection were tested with their respective hosts. Mucin presence had a positive effect for all the C or G genotype phages (FCOV-F47, -F49, -F50, -F51, -F52, -F59, -F60, -F61, -F62, V156 or FCOV-F46, -F54 and -F55, respectively; **Supplementary Fig. 3b**). It is important to note that all the phages were tested under the same conditions, and adjusting the protocols regarding incubation times, mucin concentrations and removal of biofilms to suit each would probably optimise phage growth.

### Changes in bacterial physiology elicited by mucin exposure and consequent improved phage growth is not limited to *F. columnare*

After verifying that mucin exposure changes *F. columnare* virulence and affects phage growth, we tested other bacterial species to determine whether this phenomenon is exclusive to *F. columnare*. When testing two *Aeromonas* sp. strains (B135 and B158) from our freshwater bacterial collection, we discovered that mucin exposure led to the formation of biofilm, suggesting that mucin exposure also modifies the physiology of these bacteria. Phage V46 was capable of growing better in *Aeromonas* sp. strain B135 exposed to primary fish mucus in a pattern similar to that described for *F. columnare* phages at the higher mucus dose (*p* = 0.03 at 24 h and *p* = 0.000002 at 48 h; **Fig. 5a**). Although no significant growth of V46 was initially seen in the purified porcine mucin cultures (**Fig. 5b**), changing the timing of mucin exposure from growing the cells in mucin 20 h before infection to adding mucin at the time of infection also resulted in improved phage growth (*p* = 0.03 at 24 h and *p* = 0.00006 at 48 h; **Fig. 5c**). We also tested V61 infections on *Aeromonas* sp. strain B158 in purified mucin cultures, but although a clear pattern was seen regarding higher phage growth in cells exposed to mucin, phage quantification was not possible due to difficulties in counting the plaques. These data show that mucin-mediated changes occurred similarly in at least two unrelated bacterium–phage pairs, and this may be widespread in nature among bacterial species that live on mucosal surfaces or use these surfaces as a means of finding and infecting animals.

**Figure 5:**
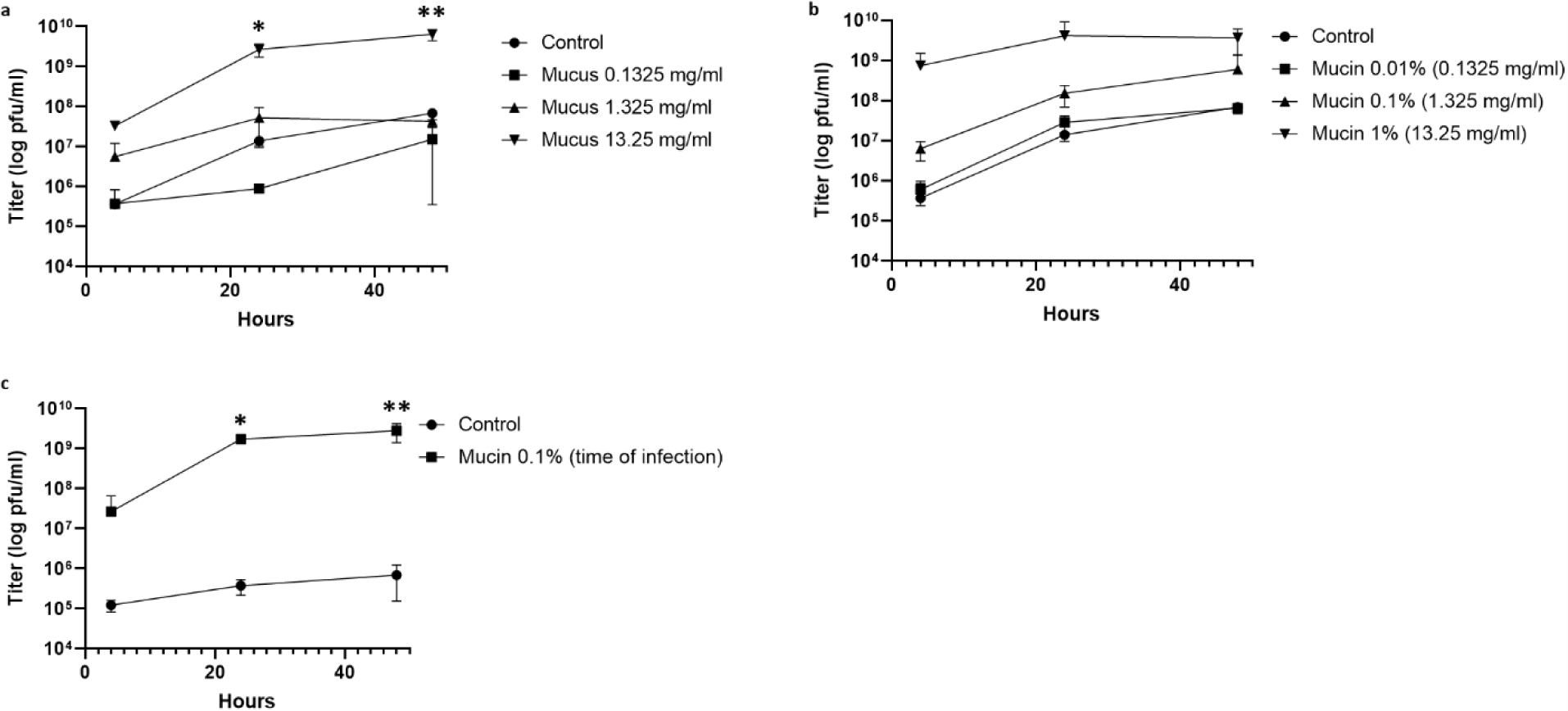
Exposure to simulated mucosal environments makes *Aeromonas* sp. strain B135 susceptible to phage V46. (**a**) V46 titre over time on 0.2x LB cultures supplemented with primary rainbow trout mucus. (**b**) V46 titre over time on 0.2x LB cultures supplemented with purified porcine mucin. For **a** and **b**, the bacteria were grown in mucus or mucin 20 h before infection with phages. (**c**) V46 titre over time on 0.2x LB cultures supplemented with purified porcine mucin at the time of infection. Each data point represents the average of triplicates and their standard deviation. Unpaired *t*-tests were used for comparing controls and tested conditions (\**p* < 0.05, \*\**p* < 0.001).

No changes in bacterial culture phenotype were seen in mucin-containing cultures of *Aeromonas salmonicida, Escherichia coli, Pseudomonas fluorescens, Salmonella enterica* or *Yersinia ruckeri* or in the Flavobacteriaceae species *Chryseobacterium indologenes, Flavobacterium* sp., *F. johnsoniae* or *F. psychrophilum.* When compared with controls, exposing hosts to mucin did not have any positive effect on the phages T4, PRD1, FL-1 or FLiP growth. The hosts used were *E. coli* DSM613, *S. enterica* DS88, *Flavobacterium sp.* B183 and *Flavobacterium sp.* strain B330, respectively (**Supplementary Fig. 4a–4d)**. Interestingly, at 0.1%, mucin had a negative effect on the growth of phages FPV4 and FPV9 on *F. psychrophilum* strain 950106-1/1 (**Supplementary Fig. 4e** and **4f**), a finding that may reveal new aspects of mucus–bacterium–phage interactions if investigated further.

Finally, we tested the ability of different phages to bind to mucin-containing agar plates. FCL-2 bound preferentially to mucin plates (*p* = 0.000001, *F. columnare* host), as did T4 (*p* = 0.000005, *E. coli* host), V46 (*p* = 0.001, *Aeromonas* sp. B135 host), FL-1 (*p* = 0.001, *Flavobacterium* sp. host) and PRD1 (*p* = 0.01, *S. enterica* host), while FLiP (*Flavobacterium* sp. host) did not (**Supplementary Fig. 4**). Except PRD1, all the mucin binding viruses were tailed phages, so Ig-like domains could be the mechanism behind the mucin interaction. PRD1’s interaction with mucin must be better investigated to confirm this finding. Due to the small plaque size and turbidity of the mucin-containing plates, it was not possible to test FPV4 or FPV9 (*F. psychrophilum* phages) or V61 (*Aeromonas sp.* B158 phage) in this experimental system. That T4 and FL-1 are also mucin-binding phages is interesting and may point to the existence of host strains that are modified by mucus exposure.

## Discussion

Interactions with animal mucus may increase the fitness of phages by limiting their diffusion in the environment, thereby increasing the probability of encountering bacterial hosts on mucosal layers. This phenomenon may be especially beneficial for metazoans if the mucus-associated phages infect pathogenic bacteria. However, how widespread the capacity for mucus binding is and how it affects the interactions between phage and bacteria are issues that remain poorly understood. Inspired by the BAM model proposed by **Barr et al.**^**8**^ and using a natural infection system consisting of rainbow trout, its bacterial pathogen *F. columnare* and phage FCL-2 (capable of infecting the bacterium), we studied how the mucus environment affects phage–bacterium interactions. We showed that an *F. columnare*–infecting phage binds to primary fish mucus and purified porcine mucin at comparable levels to those of phage T4. After a single exposure of fishes to FCL-2, the phage was held for up to 1 week on the skin mucosa. When fish were simultaneously exposed to FCL-2 and T4, both phages were recovered from mucus with similar titres for up to 1 week, showing that the phages do not interact antagonistically in mucus even at high densities.

The interaction between mucins and phages has been suggested to occur via Ig-like domains in the phage capsid. The orfs in FCL-2 do not show any similarity to the highly antigenic outer capsid protein (hoc) of T4, which mediates adherence to mucus^**8**^. However, FCL-2 orf17 has homology with a phage protein possessing an Ig-like fold, the lactococcal phage TP901-1 baseplate wedge protein. This implies that the mechanisms for persisting in the mucus for T4 and FCL-2 may be similar, and the sub-diffusive motion created by phage Ig-like folds weakly interacting with mucins creates a vector of movement that is stronger than the mucus shedding and water flow. In addition, the *Aeromonas* sp. podovirus V46, which was also enriched in mucin-containing plates, has a highly similar capsid protein to that of phage P22, and both phages possess the telokin-like Ig domain in the capsid protein. The mucin binding mechanism of the coliphage PRD1, the type of member of Tectiviridae, remains unclear because it does not have distinct Ig-like folds in its structural proteins. However, even weak folds multiply in the capsid and could mediate mucin interaction.

Expanding on the theme of phage mucus adherence and protection of metazoans from infections, our data demonstrate that the prophylactic use of phages in phage therapy can be relevant and effective. Phage concentrations remained relatively high in the rainbow trout mucus for 1–3 days after exposure, despite constant water flow. In addition, these adhered phages provided protection against bacterial infection. We observed a significant delay in the onset of disease and increase in fish survival, although complete protection was not seen. It is important to note that the conditions of the experiment with the high dose of bacteria used favoured the bacterial infection and not the phage. Still, even the very low number of phage particles observed in mucus (one individual fish would contain less than 200 FCL-2 particles) was enough to hold back the infection resulting from exposure to 9.96 × 10^8^ cfu of *F. columnare* (equivalent to using an approximated multiplicity of infection of 1 × 10^−6^ in each tank). In nature or at fish farms, the number of infectious bacteria is likely to be lower, making protection by mucus-bound phages even more relevant. In phage therapy, the use of phage cocktails reduces the chances of selecting for resistant bacteria, but the mucus-binding capacities of different phages may vary. Interestingly, our results also imply that the co-occurrence with ‘non-relevant’ phages may influence protection against infection. Our data suggest that pre-exposure to FCL-2 combined with T4 protects fish longer than FCL-2 alone, although T4 did not replicate in the system. The mechanism behind this finding is still not known, but since the T4 particle is larger than that of FCL-2, it may help in keeping FCL-2 in the lower mucus layer. More studies are needed to understand how mucus influence the interactions between phages and how combinations of different phages (phage cocktails) may influence protection against infection.

Exposing metazoans to phages as prophylaxis against bacterial infections has been tested in the past. In 2017, a cocktail of three phages against *Vibrio cholerae* was shown to be effective if given orally 24 hours before infecting mice and rabbits, and phage retention on the intestine was shown^**24**^. In another study, a phage infecting *Salmonella enterica* serovar Enteritidis was more efficient in protecting poultry if given prophylactically by an oral route. Phages were recovered from faeces 7 days after treatment, even in the group not challenged with the bacterial host^**25**^. Furthermore, massive human trials based on preventive phage therapy (called ‘prophylactic phaging’) were performed in the past by the former Soviet Union. Phage pre-treatment was noted to reduce disease occurrence, and phage persistence for days in the human body was recorded^**26**^. Although the exact mechanism of phage retention was not studied, it is possible that the success of prophylactic use of phages, in experimental conditions and human trials, is mediated at least in part by phage sub-diffusion on mucosal surfaces. Sub-diffusive motion in mucus would hold phages on the mucosa, allowing the phages to find and infect invading mucosal pathogens. For non-mucosal bacterial infections, the mucus could still retain and shed phages over time, resulting in prolonged protection. The interaction of phages and metazoans may be even more profound, since phages have been shown to pass through animal cells^**27**^. On a mucosal surface, sub-diffusive motion towards the mucus source could serve as a gateway to transcytosis, making phages approach and cross apical cell membranes. The efficiency of phages in protecting mucosae contrasts with a recent finding that mucus layers may negatively affect antibiotic activity on *Pseudomonas aeruginosa*^**28**^. This highlights the relevance of considering phages as a treatment for mucosal infections.

In addition to phage retention by mucus and prophylactic use against *F. columnare*, we investigated bacterium and phage interactions on simulated mucosal environments. Culture media containing primary mucus from rainbow trout or purified porcine mucin affected the bacterial characteristics and phage growth. More recently, mucus exposure was shown to activate *F. columnare* biofilm formation and regulate transcription^**29**^, illustrating that mucins are not only used as nutrients but also function as transcriptional regulators. In contrast, transcriptional upregulation of mucin genes has been associated with catfish susceptibility to *F. columnare*^**30**^. Our results add to this knowledge by showing that exposure to mucin can also directly increase bacterial virulence. Mucin elicited changes in growth in all the *F. columnare* strains tested but not in other Flavobacteriaceae species. It can be taken as an indication that the close relationship of *F. columnare* with its host mucus, demonstrated by the disease characteristics, is the major driver of the selection of the mucus-induced changes. As such, mucin exposure upregulation of bacterial virulence may not be exclusive to *F. columnare*. We found two *Aeromonas* sp. strains that also form biofilm as a response to primary mucus or mucin exposure. In addition, it has been described that mucin exposure increases the mucus-binding capacity of *Lactobacillus reuteri* strains, and in this case, mucin may also be acting as a signal that changes cell transcriptional status^**31**^.

Our data add another component to the interaction between bacteria and mucus by considering phages and the BAM model. The results indicate that the mucin-induced upregulation of *F. columnare* virulence has trade offs in increasing bacterial susceptibility to phages. Mucus-adhering phages may have evolved to exploit the mucus-induced upregulation of bacterial virulence–related factors for infection. This emphasises the ecological relevance of microbial interactions occurring in the eukaryotic mucus environments, with implications for the outcome of columnaris disease. In the case of *F. columnare*, the possible mechanism causing this comprise both the increased bacterial growth and expression of flavobacterial gliding motility in presence of mucus, which may increase the availability of surface receptors for phage infections. Colony spreading is used as a measure of flavobacterial gliding motility, which is linked with the type IX secretion system of virulence factors^**32**^. Loss of gliding motility has been associated with a decrease in bacterial virulence and increase in phage resistance for *F. columnare*^**33**^. Nevertheless, the mucus-induced changes in bacterial phage susceptibility enabled to overcome the decades-old difficulties regarding production of *F. columnare* phages in liquid cultures^**34**^. This opens possibilities for isolating new *F. columnare*–associated viruses and growing them for large-scale purposes. Further research is needed to resolve how mucus components affect the phage–bacterium interactions in other mucus-related bacterial pathogens.

Mucus may have a significant effect for phage–bacterium interactions on a wider scale. A study focussing on *Clostridium difficile* and phage interaction on human cell cultures concluded that the phage more efficiently reduces the number of planktonic and adherent bacterial cells in the presence of the human cells than occurs *in vitro*^**35**^. Although the hypothesis for this finding was that increased phage activity was related to strong phage adsorption to the cells, no data were presented regarding changes elicited by the human cells’ effect on the bacterium. The HT-29 cells used are known to be heterogeneous, containing a small percentage of mucin secreting cells, and HeLa cells produces at least MUC1^**36,37**^, so the possibility exists that the released cell-derived mucus changes the bacterial physiology and increases its susceptibility to phages. Comparison of three *Vibrio cholerae* phages’ predation in different niches revealed that two are only capable of infecting in nutrient-rich conditions^**38**^, which could also imply an importance of mucin for changing the bacterial host cells. When testing our collection of freshwater bacteria, we found two *Aeromonas* sp. strains that behaved similarly to *F. columnare* in terms of biofilm formation and phage susceptibility after primary mucus and purified mucin exposure. Although no virulence data exist for these isolates, the *Aeromonas* genus contains many putative disease-causing species^**39**^, so interaction with metazoan mucus may also play an important role for these organisms and its interaction with phages.

Our data confirm that phage adhesion to mucus is important for metazoan protection, provides a system in which the BAM model can be tested and shows that investigating phage–bacteria interactions in mucus is relevant for mucosal pathogens. It also provides important information regarding the mechanism behind the use of phages for prevention and highlights the relevance of the use of ‘phaging’ in future phage therapy trials. By preventively enriching the metazoan mucus layers with added phages, it would be possible to modulate the mucus microbiome, thereby generating a protective phage layer that could help to avoid disease instead of only treating it as done in conventional phage therapy. Thus, a prophylactic phage therapy approach can have a positive effect on public health and protecting farmed animals worldwide. In conclusion, our model has implications that go beyond columnaris disease and can be applied to better understand phage–bacterium interaction on metazoan mucus in a broader sense.

## Methods

### Bacteria and phages

The bacterial strains and phages used in this study are listed in **Table 1**. *F. columnare* strain B185 and its phage FCL-2 (myophage) were chosen as models because they have been well characterised and shown to be relevant for phage therapy studies^**21**^. The other strains were used for validation of our observations.

### Growth conditions, phage titrations, primary mucus and mucin preparation

*F. columnare, F. johnsoniae, Chryseobacterium indologenes* and *Flavobacterium* sp. strains were cultured in modified Shieh medium (without glucose, as described by **Song et al. 1988**)^**40**^ at 25°C and 120 rpm. *F. psychrophilum* was cultivated in TYES media^**41**^, at 15°C and 120 rpm. *E. coli* and *S. enterica* were cultivated in LB medium at 37°C and 200 rpm. *Aeromonas salmonicida, Aeromonas* sp. strains, *Pseudomonas fluorescens* and *Yersinia ruckeri* were cultivated in 0.2x LB medium at 25°C and 120 rpm. The phages were titrated using the double-layer agar method on the appropriated hosts. Bacterial inocula were normalised by adjusting the OD595 from overnight cultures. In the case of *F. columnare*, a regression curve was used for estimating colony forming units (cfu) from OD values to calculate the inocula and multiplicity of infection (moi) whenever necessary.

Primary mucus was collected from rainbow trout individuals (*Oncorhynchus mykiss*). Fish (mean size of 8 cm) were euthanised with an overdose of benzocaine, and primary mucus was harvested by scraping the skin with a glass slide. Mucus was pooled on Falcon tubes on ice and briefly centrifuged (1000g, 5 min, 4°C) to remove scales and other debris; then, the supernatant was autoclaved and the sterile mucus was aliquoted and stored at –20°C until use. The total protein concentration was determine by nanodrop measurements. As an example, a batch made from 277 fish yielded around 100 ml of primary mucus with a total protein concentration of 26.5 mg/ml. Purified porcine mucin (Sigma, CAT# M1778) was diluted in water to a 2% final concentration, autoclaved and stored at +4°C. The total protein concentration of a 2% mucin solution varied from 20 to 26 mg/ml depending on the batch.

Simulated mucosal cultures were prepared by mixing complete culture media to an equal volume of primary mucus or purified porcine mucin and water, resulting in 0.5× media containing the desired amount of mucus or mucin. Doses of 0.1325, 1.325 and 13.25 mg ml^-1^ of primary mucus were used, which corresponded to the total protein concentration of 0.01, 0.1 and 1% purified mucin solutions respectively. Controls consisted of 0.5× media. Optimal results for *F. columnare* were obtained by inoculating five milliliters of media with 5×10^4^ cfu, and then infecting the cultures 22 hours later with phages (moi 0.01, estimated from the control bacterial number).

### Identification of the Ig-like domain in phage genomes

The genome of phage FCL-2 has been sequenced^**21**^. Here, the genome was re-annotated (September 2018) using protein BLAST^**42**^ and HHPred^**43,44**^ for protein homology detection and structure prediction. HHPred was also utilised for analysing orfs from *F. psychrophilum* phages FpV-4 and FpV-8, *Salmonella* phage PRD1 and the capsid protein of *Aeromonas* sp. phage V46. In addition, Phyre2^**45**^ was used for protein fold recognition of predicted structural proteins.

### Phage retention by rainbow trout skin mucus

Thirty fish (mean size 8 ± 0.96 cm) were incubated with 1.25 × 10^12^ pfu of FCL-2 in 2 L of water (6.25 × 10^8^ pfu ml^-1^) for 30 min. Then, 4 L of fresh water was added into the aquarium. Three hours later, the water flow was opened, resulting in a change of water at a rate of 860 ml min^−1^ over the duration of the experiment. Aerated water was maintained at 15°C, and the fish were fed daily. At the indicated timepoints, three fishes were collected from the aquarium and euthanised with benzocaine; their skin mucus was scraped with the help of a glass slide and mixed with 450 µl of Shieh medium. Three independent water samples (450 µl each) were also collected. All the samples were preserved by the addition of chloroform (10% final concentration) and used later for phage titrations.

### Prophylactic phage therapy experiment

One week before the experiment, 118 rainbow trout (mean size 8.3 ± 1.12 cm) were moved from the stock tanks to experimental aquaria and acclimatised to 25°C (an increment of +2°C per day, except on the first and last days). At 7, 3 and 1 days before the infection, designated groups of fishes were exposed to phages. Exposure was made by moving the fish to 4 L of water containing 1 × 10^12^ pfu of FCL-2 (2.5 × 10^8^ pfu ml^−1^) alone or a mixture of 1. × 10^12^pfu of FCL-2 and 1.2 × 10^12^ pfu of T4 in 4 L of water (2.5 × 10^8^and 3 × 10^8^ pfu ml^−1^, respectively). One aquarium was used for each condition: five for the phage pre-treatments and two for controls (fishes not exposed to phages). After 30 min of exposure to phages, the water flow was opened (water flow was 248 ± 35 ml/min). The fish were fed daily until the time of infection and kept in aerated water during all the experiments.

On the day of infection, the fish were moved to aquaria containing 2 L of water (each of the seven original aquaria was divided into three, with five fish in each), with no flow through. Before transferring the fish from the phage treated groups, they were immersed in freshwater for minimising the carryover of phages not attached to the mucus. Then, the bacteria were added to these aquaria for the bacterial immersion challenge (see below), and 2 h later, the fishes were moved to larger aquaria containing 6 L of freshwater (no flow through, divisions of three aquaria per tested condition kept). The fish were checked regularly, and dead or moribund fish were removed from the aquaria.

The bacterial inoculum was prepared by adding 2 × 10^6^ cfu of *F. columnare* strain B185 to 200 ml of Shieh (triplicates). Twenty-four hours later, at the time of infection, the bacterial number was estimated by OD measurements and confirmed by titrations. The density of bacteria used for infections was 9.96 × 10^8^ pfu per aquarium (4.98 × 10^5^ cfu ml^-1^ in the 2 L used for the immersion challenge).

The mucus of three fish from each condition tested (phage pre-treatments and controls) was collected before the infection process. Water from the original aquaria (where the fishes and phages were kept) was also collected at the same time. At 10, 20, 34 and 48 h after infection, water from all the aquaria was collected. All the samples were preserved by the addition of chloroform to a final concentration of 10% and used for phage titrations and qPCR.

### Quantitative PCR

To quantify the *F. columnare* genome copies from aquarium water, a quantitative PCR reaction was designed to detect the CRISPR spacer region in strain B185. The reactions were performed with 5 μl of water samples as template, 0.3 µM of forward (ATTTGGCGGTTGACCATAGAT) and reverse (CGGTGTCCACTTGAAATACCTTAC) primers, and 10 μl of 2x SYBR Green Supermix (iQ SYBR Green Supermix, Bio-Rad) in a 20-μl reaction volume. All the samples were amplified in triplicate. The PCR reaction started at 95°C for 3 min for initial denaturation, followed by 40 cycles of 15 s at 95°C for denaturation and 60 s at 60°C for annealing. High-resolution melting analysis was performed immediately afterwards by increasing the temperature from 55°C to 95°C by steps of 0.5°C maintained for 5 s each. A standard curve for the quantification of B185 was generated by analysing a 10-fold dilution series of its DNA with known quantities of 5 ng μl^−1^ to 0.005 pg μl^−1^ 1 (equivalent to 2 × 10^6^ to two copies).

### Helium ion microscopy

Biofilm samples were prepared by adding a sterile 3 × 3 mm piece of agar to 5 ml of 0.5× Shieh supplemented with 0.1% mucin culture and then inoculating the culture with 5 × 10^4^ cfu of *F. columnare* strain B185. On the next day, the agar piece colonised by the biofilm was fixated and dehydrated, as described in **Leppanen et al.** ^**46**^ prior to imaging in a helium ion microscope (Zeiss).

### Chemotaxis assay

The cell density of overnight cultures of *F. columnare* strain B185 was adjusted to 1 × 10^8^ cfu ml^−1^ and 200 µl was added to the wells of a 96-well plate. By using a multichannel pipette (Thermo Scientific, CAT# 46300500), 200 µl of the chemoattractants (consisting of 0.5× Shieh alone or supplemented with mucin) were kept inside the pipette tips and inserted into the wells containing the bacteria. After 90 min, the tip contents were carefully dispensed into clean wells, serially diluted and titrated. The relative chemotaxis response (RCR) was calculated by dividing the cfu count of the tested conditions by the cfu count of the control (0.5× Shieh).

### Spread on agar and protease secretion

The cell density of overnight cultures of *F. columnare* strain B185 was adjusted to 1 × 10^6^ cfu ml^−1^, and 10-µl drops were added to the top of Shieh-agar plates supplemented or not with mucin and 1.5% skimmed milk. The colony and protease halo sizes were measured over time in millimetres.

### Virulence *in vivo*

The experimental setting was similar to that of the prophylactic testing described above but using 59 fish (mean size 8.08 ± 1.17 cm) and no phage treatments. The bacterial inoculum was prepared by adding 2 × 10^6^ cfu of *F. columnare* strain B185 to control cultures (0.5× Shieh) or 0.5× Shieh supplemented with 0.1% mucin cultures 1 day before the infection. For each condition, three individual 200-ml cultures were prepared. Twenty-four hours later, at the time of infection, the bacterial number was estimated by OD measurements and confirmed by titrations. In the case of mucin cultures, only the supernatant (containing planktonic cells) was used, and the biofilms were discarded. The doses used for infection were: 2.17 × 10^6^ and 4.22 × 10^5^ cfu ml^−1^ for controls and 1.83 × 10^5^ cfu ml^−1^ for mucin-grown bacteria in the 2 L of the immersion challenge. The fish were monitored frequently for disease symptoms and morbidity. Morbid fish that did not respond to external stimuli were considered dead and removed from the experiment.

### Testing the susceptibility of biofilms and planktonic cells to FCL-2

*F. columnare* B185 mucin cultures (0.1%) were prepared as described above. Before infection with FCL-2, the cultures were either kept as a whole (biofilm and plankton together, as in the previous experiments) or had the supernatant containing planktonic cells transferred to a new flask. Five millilitres of fresh Shieh medium were added to the biofilms left without plankton. All the cultures were infected with the same amount of virus (moi 0.01 based on the cfu estimative of the controls), and the phages were quantified by titration after 4 and 24 h.

### Evaluating the effect of FCL-2 on the *F. columnare* biofilm spread

*F. columnare* B185 was added to 0.5× Shieh supplemented with 0.1% mucin containing sterile plastic biofilm baits (used in recirculating aquaculture systems). On the next day, one piece of colonised plastic was transferred to fresh 0.5× Shieh supplemented with 0.1% mucin media, containing more sterile biofilm baits, in the presence or absence of FCL-2. The results were assessed qualitatively by observing the amount of biofilm formed on the flask walls and the sterile biofilm baits 24 h later.

### Testing the duration of the mucin effect

*F. columnare* strain B185 was grown in control (0.5× Shieh) and 0.5× Shieh supplemented with 0.1% mucin cultures. Six cultures were made for each condition, and these cultures were considered to be passage 0. On the next day, 5 µl of each culture was transferred to fresh 0.5× Shieh, and this was called passage 1. Then, half of each of the original tubes (from passage 0) was infected with phage FCL-2. On the next day, the same process was repeated with passage 1 to create passage 2, followed by infection of half the tubes in passage 1. Phages and bacteria from every passage were titrated for 24 h after phage infection. The whole process was repeated until passage 3.

### Phage retention by purified porcine mucin

Five millilitres of media containing phages were added to the top of agar plates supplemented with 1% mucin or without mucin as control. Concentrations of 2.5 × 10^3^ pfu ml^−1^ were used for PRD1, FCL-2, V46, FLiP and FL-1. For T4, 2.5 × 10^2^ pfu ml^−1^ was used due to large plaque sizes. After 30 min of shaking at room temperature, the liquid was removed; the plates were left to dry for 30 min, and then 3 ml of soft agar containing 300 µl of an overnight culture of the bacterial host was added to each plate. The plates were incubated at room temperature, and the plaques were counted 2–3 days later to measure the amount of phages held on the plates.

### Statistical analysis

Analysis was done using the GraphPad software version 8.0.1. The log-rank (Mantel– Cox) test was used for evaluating the survival curves. Unpaired *t*-tests were employed for comparing controls and tested conditions in the appropriate datasets.

### Animal experimentation

Fish experiments were conducted according to the Finnish Act on the Use of Animals for Experimental Purposes, under permission ESAVI/3940/04.10.07/2015 granted for Lotta-Riina Sundberg by the National Animal Experiment Board at the Regional State Administrative Agency for Southern Finland.

## Acknowledgments

We would like to thank Professor Jean-Francois Bernardet (INRA, France), Professor Mark McBride (University of Wisconsin—Milwaukee, USA) and Professor Mathias Middelboe (University of Copenhagen, Denmark) for kindly donating bacterial and phage strains (**Table 1**), as well as Jonatan Pirhonen for technical help. This work was supported by the Finnish Centre of Excellence Program of the Academy of Finland (CoE in Biological Interactions 2012-2017 #252411), Academy of Finland (grants #266879 and #304615) and Jane and Aatos Erkko Foundation. This work resulted from the BONUS FLAVOPHAGE project supported by BONUS (Art 185), funded jointly by the EU and Academy of Finland.

## Author contributions

G.M.F.A. and L.R.S. designed the experiments. G.M.F.A. performed the experiments. E.L. was responsible for bioinformatics and the collection of phage and bacterial strains. R.A. was responsible for designing the quantitative PCR and assisting on sequencing for bacterial identification. All authors contributed to writing the manuscript.

---

**Supplementary Figure 1:**
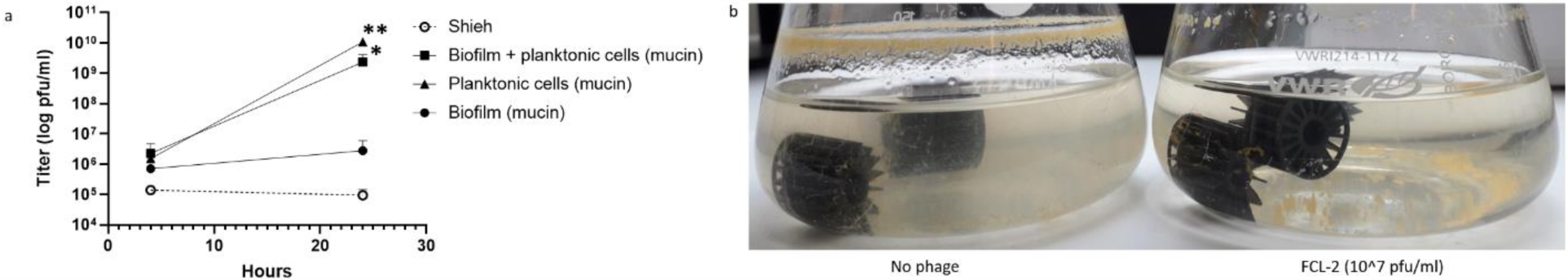
Phage FCL-2 infects planktonic cells from simulated mucosal cultures and limits biofilm spread. (**a**) FCL-2 titres on 0.5× Shieh cultures or in 0.5× Shieh cultures supplemented with 0.1% mucin and infected either as a whole (biofilm and plankton together) or divided into biofilm and planktonic cell portions. (**b**) Qualitative evaluation of the effect of FCL-2 presence on the biofilm spread in mucin-containing cultures. Each data point represents the average of triplicates and their standard deviation in **a**. Unpaired *t*-tests were used for comparing controls and tested conditions (\**p* < 0.05, \*\**p* < 0.001).

**Supplementary Figure 2:**
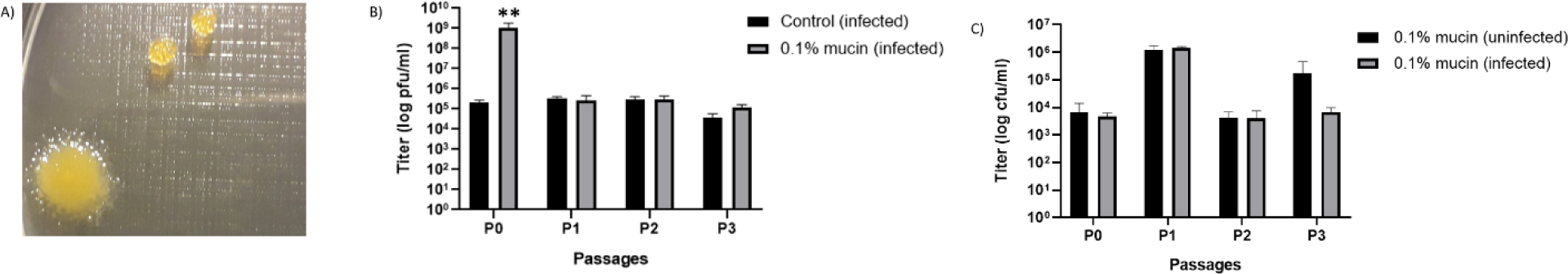
Phage susceptibility induced by simulated mucosal culture exposure disappears faster than morphological colony changes do. (**a**) Aspect of a new colony morphotype appearing on mucin-containing cultures of *Flavobacterium columnare* (left). Two conventional rhizoid colonies are evident in the top right. (**b**) FCL-2 phage titres on sequential passages of 0.5× Shieh and 0.5× Shieh supplemented with 0.1% mucin cultures. (**c**) Colony-forming units of *F. columnare*–modified rhizoid colonies from sequential passages of 0.5× Shieh and 0.5× Shieh supplemented with 0.1% mucin cultures. Each data point represents the average of triplicates and their standard deviation. Unpaired *t*-tests were used for comparing controls and tested conditions (\*\**p* < 0.001).

**Supplementary Figure 3:**
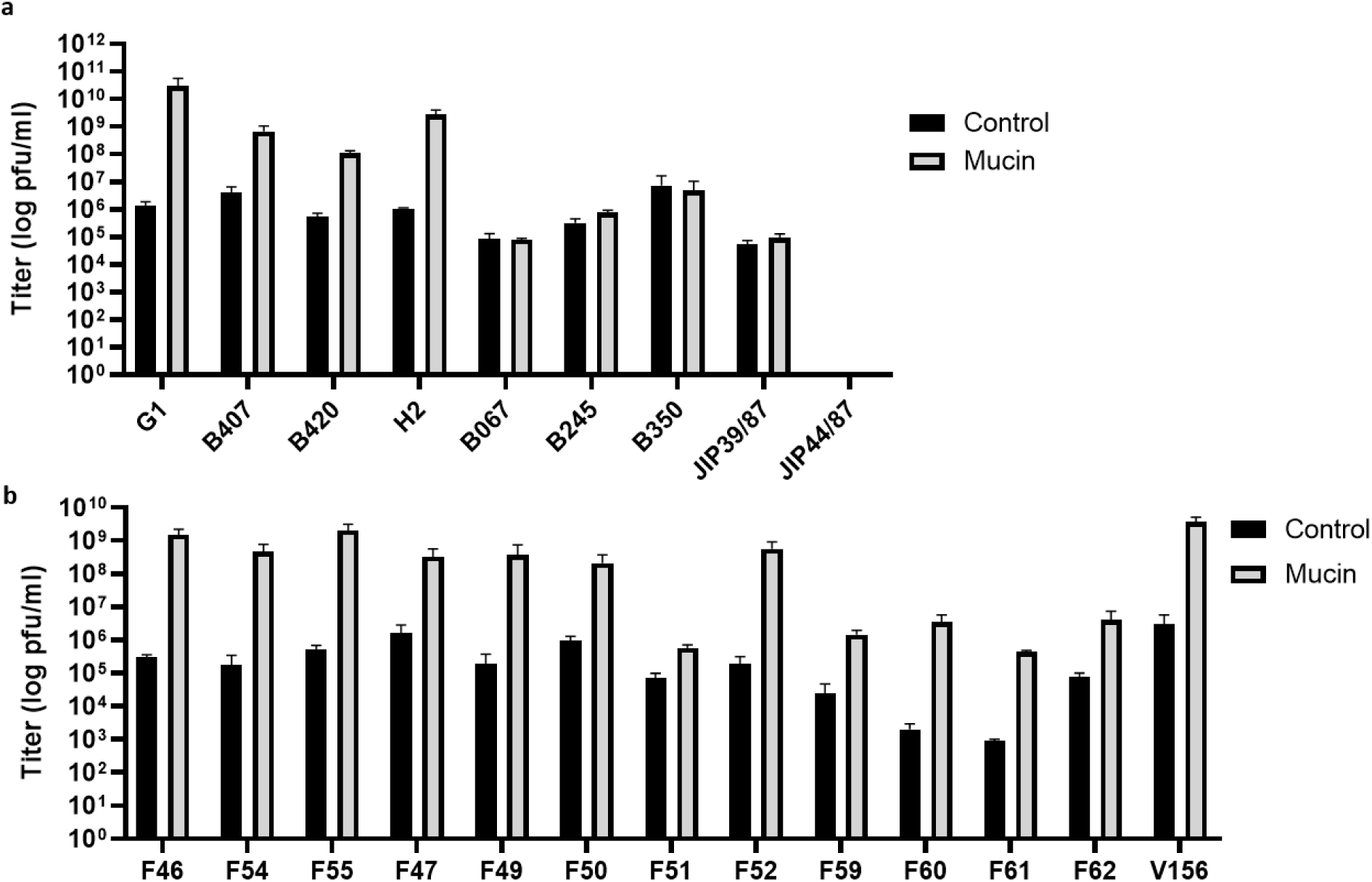
Effect of exposing *Flavobacterium columnare* to mucin for phage growth. (**a**) Phage FCL-2 titres after infections in different *F. columnare* strains grown in 0.5× Shieh alone or supplemented with 0.1% mucin. (**b**) Different *F. columnare* phage (FVOV-F46-62, V156) titres after infection on their respective hosts grown in 0.5× Shieh alone or supplemented with 0.1% mucin. Each data point represents the average of triplicates and their standard deviation, except B245 and B350 in **a** and V156 in **b**, which were done in duplicate.

**Supplementary Figure 4:**
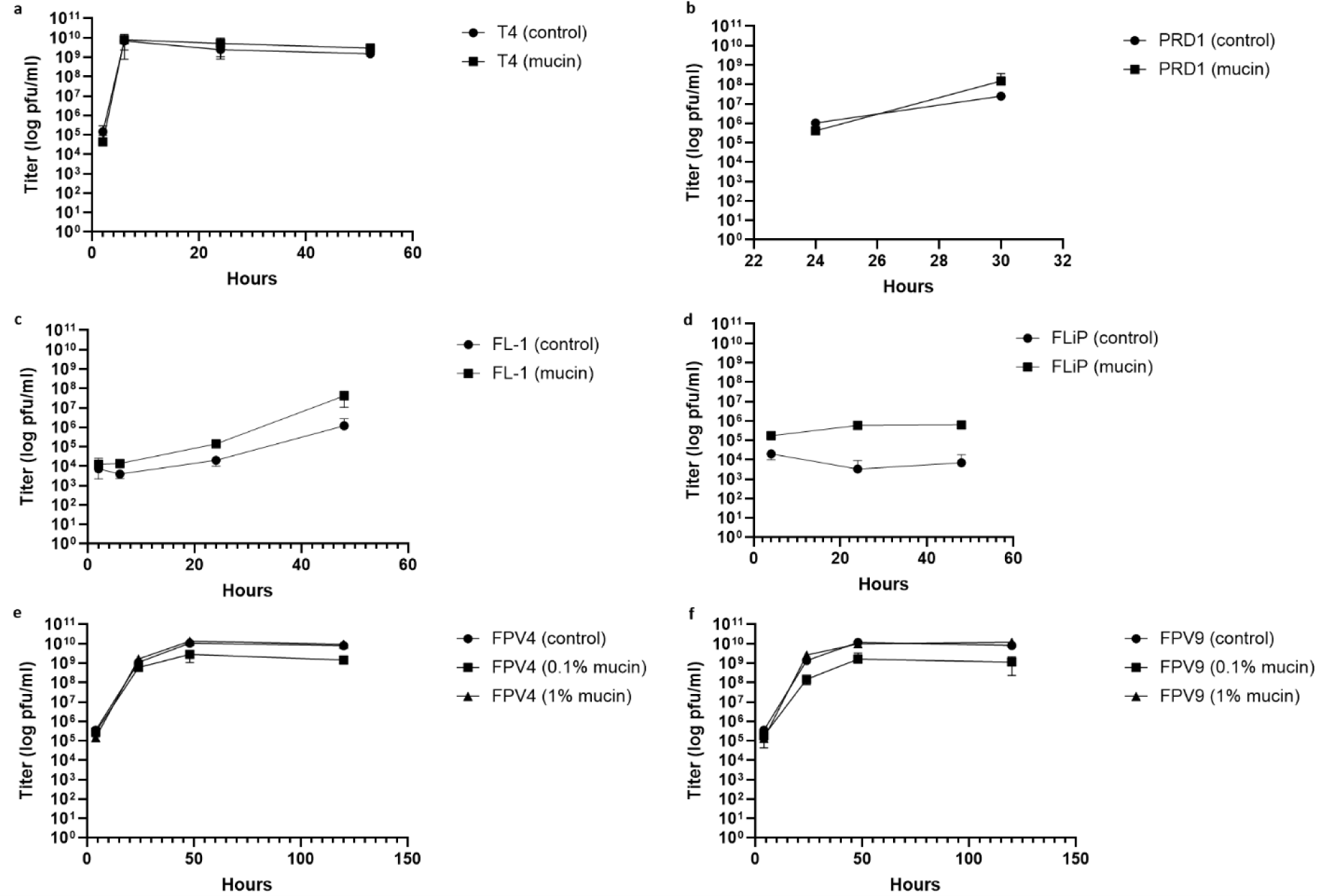
Effects of mucin exposure on phage–host interactions in different bacterium–phage pairs. (**a**) T4 titres after infection in *E. coli* grown in 0.5× LB alone or supplemented with 1% mucin. (**b**) PRD1 titres after infection in *S. enterica* grown in 0.5× LB alone or supplemented with 1% mucin. (**c**) FL-1 titres after infection in *Flavobacterium* sp. grown in 0.5× Shieh alone or supplemented with 1% mucin. (**d**) FLiP titres after infection in *F.* sp. grown in 0.5× Shieh alone or supplemented with 1% mucin. (**e**) FPV4 titres after infection in *F. psychrophilum* grown in 0.5× TYES alone or supplemented with mucin. (**f**) FPV9 titres after infection in F. *psychrophilum* grown in 0.5× TYES alone or supplemented with mucin. Each data point represents the average of triplicates and their standard deviation, except **e** and **f**, which were performed in duplicate and confirmed in an independent experiment.

**Supplementary Figure 5:**
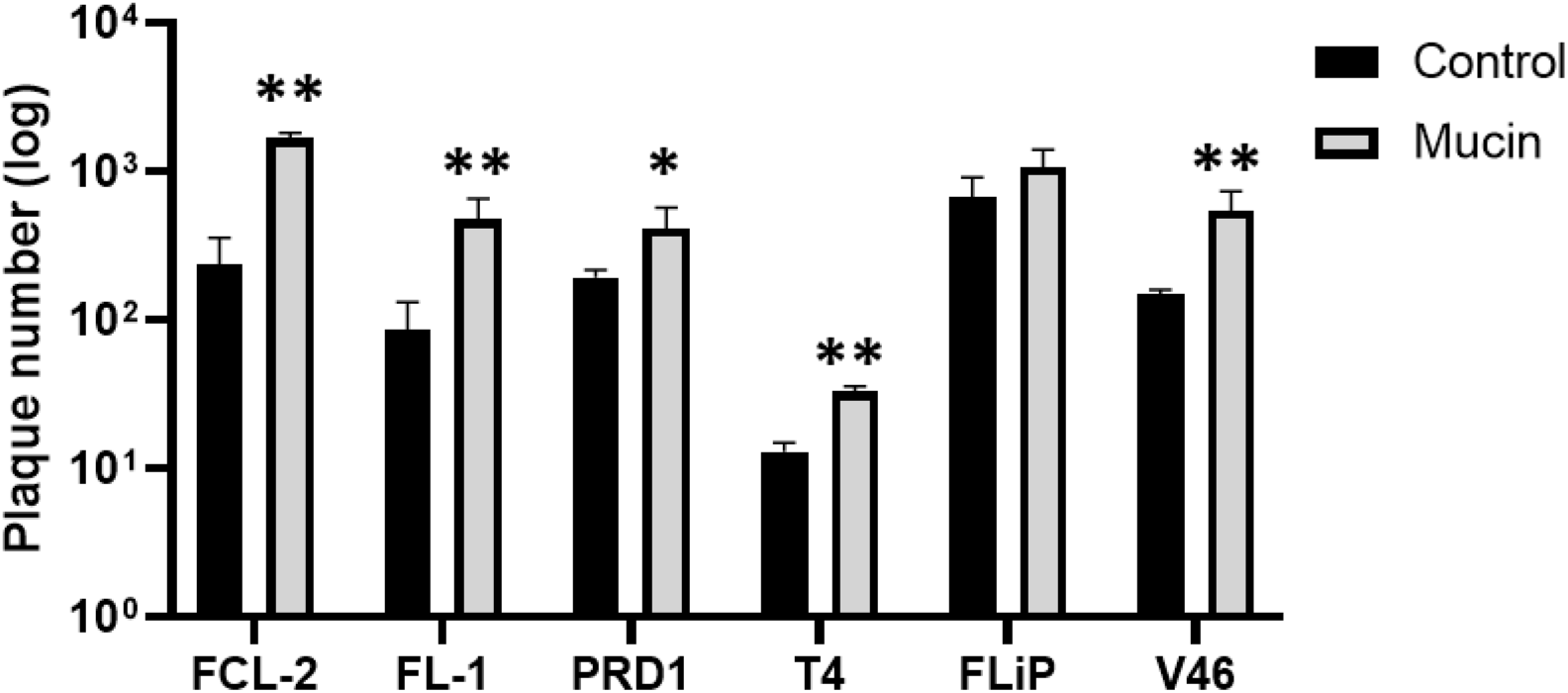
Phage adhesion to mucin-containing agar plates. Bars represent the amount of plaques counted in the control agar plates or agar plates supplemented with 1% purified porcine mucin. Each data point represents the average of triplicates and their standard deviation. Unpaired *t*-tests were used for comparing controls and tested conditions (\**p* < 0.05, \*\**p* < 0.001).

